# Ab-VS: Evaluating Large Language Models for Virtual Antibody Screening via Antibody-Antigen Interaction Prediction

**DOI:** 10.1101/2025.07.26.666985

**Authors:** Yiming Zhang, Koji Tsuda

## Abstract

We present Ab-VS-Bench, a new benchmark for evaluating large language models (LLMs) on antibody virtual screening (VS) tasks through natural language instructions. Unlike prior efforts that focus on structural annotation, binding prediction, or developability using frozen antibody models, Ab-VS-Bench targets the end-to-end VS workflow and leverages LLMs’ instruction-following capabilities. The benchmark comprises three core tasks inspired by small-molecule VS: (1) *Scoring*—predicting antibody–antigen binding affinity, (2) *Ranking*—ordering antibodies by affinity or thermostability, and (3) *Screening*—identifying high-affinity binders from large antibody libraries. We convert experimental datasets into instruction-tuning format and evaluate multiple model variants, including zero-shot LLMs, instruction-finetuned LLMs, and multimodal models enhanced with antibody-specific embeddings. Ab-VS-Bench provides a unified framework to benchmark LLMs for antibody discovery, aiming to catalyze progress in language-guided biotherapeutic design.

## I. Introduction

Antibodies are critical biomolecules widely used in therapeutics and diagnostics due to their high specificity and binding affinity to diverse antigens [1]. Designing or discovering antibodies with favorable binding and biophysical properties is a central goal in drug development [2]. In small-molecule drug discovery, *virtual screening*—a computational process that filters large chemical libraries—has significantly accelerated hit discovery [3]. A similar approach, if applied to antibody discovery, could streamline the identification of potent binders from a vast combinatorial sequence space.

However, most current computational methods for antibody screening rely on structure-based modeling or large-scale docking simulations, which are expensive and not easily scalable [2]. Meanwhile, large language models (LLMs), trained on vast corpora of natural language and biological sequences, have shown impressive instruction-following, reasoning, and generalization capabilities [4], [5]. This raises a natural question: *Can LLMs, when prompted in natural language, perform virtual antibody screening*?

To date, no benchmark has been specifically designed to test this capability [6]. Existing efforts such as NbBench [7] provide a comprehensive evaluation suite for nanobody modeling, covering eight biologically meaningful tasks including CDR annotation, binding prediction, thermostability regression, and polyreactivity classification. While valuable for assessing frozen representations of pretrained models, NbBench focuses on isolated classification or regression tasks and does not simulate the end-to-end screening workflow. It also does not assess instruction-following abilities or interactive reasoning over candidate sets—key strengths of modern LLMs.

To address this gap, we introduce **Ab-VS-Bench**^1^, the first benchmark specifically designed to evaluate how well LLMs can perform virtual screening-inspired tasks for antibodies using natural language instructions. Ab-VS-Bench mirrors the small-molecule screening pipeline and comprises three core tasks [8]:

- **Scoring** [9], [10]: Predict the binding affinity or strength between an antibody and an antigen.
- **Ranking** [2], [11]: Sort a set of candidate antibodies based on target properties such as affinity or thermal stability.
- **Screening** [2], [11]: Identify high-affinity binders from large antibody libraries.

All tasks are framed as instruction-following problems, where the model receives a textual prompt (e.g., “Rank these antibodies by their binding affinity to EGFR”) and is expected to produce human-readable outputs such as class labels, ranked lists, or candidate selections.

We further explore three modeling regimes: (1) zero-shot LLMs (e.g., Mistral [12], Qwen2.5 [13]), (2) instruction-finetuned [14] LLMs, and (3) multimodal models that integrate antibody-specific embeddings from ESM-2 [15] via cross-attention mechanisms [16].

**Ab-VS-Bench** is constructed from experimental datasets and includes standardized splits, instructions, and evaluation metrics (e.g., Accuracy, F1, Enrichment Factors). Across all tasks, we find that fine-tuning improves performance significantly, and that multimodal LLMs outperform plain LLMs, particularly in ranking and generalization settings.

In summary, Ab-VS-Bench provides a unified and reproducible platform to evaluate general-purpose and domain-specialized models on realistic antibody screening scenarios. It highlights the potential of LLMs to go beyond static prediction, enabling interactive, language-guided decision-making in antibody design.

## II. Related Work

### Benchmarks for Antibody and Nanobody Modeling

Recent work has introduced several benchmarks to evaluate language models on antibody-related tasks. One prominent example is NbBench [7], which provides a comprehensive suite for nanobody modeling across eight tasks. These include CDR loop prediction, antigen-binding classification, affinity regression, stability prediction, and developability assessments. NbBench standardized datasets from sources like SAbDab and INDI and evaluated a variety of pretrained models—including general protein LMs, antibody-specific LMs, and nanobody-specific LMs—under a frozen setting, where model weights are fixed and only lightweight classifiers are trained.

Results from NbBench showed that antibody-specific models perform best on binding classification tasks, but all models struggled with regression tasks like affinity or stability prediction. However, NbBench frames each task as a standalone supervised problem, without exploring instruction-following or multi-step reasoning. It does not evaluate whether models can interactively perform screening workflows or respond to prompts using natural language.

Another relevant benchmark is ATUE [17], which tests antibody language models on biologically meaningful tasks such as paratope prediction and somatic hypermutation analysis. While ATUE provides insights into antibody biology understanding, it also uses task-specific models and does not involve instruction-based interaction.

AbBiBench [2] is a more recent effort focused on affinity maturation. It compiles over 155,000 mutant antibody sequences across 9 antigen targets (e.g., influenza, HER2, SARS-CoV-2) and evaluates models by how well their internal scores—like sequence likelihoods or structure-based energies—correlate with experimental binding affinities. AbBiBench also explores generative design. However, it does not involve prompting or use language model outputs directly as predictions.

### Large Language Models in Biomedicine

Concurrently, large language models (LLMs) such as GPT-3 [18] and LLaMA [19] have demonstrated strong performance in biomedical applications, from answering medical questions to analyzing genomic information [20], [21]. Applying these models to non-linguistic data like protein sequences presents a unique challenge. An effective strategy is to combine specialized biomolecular encoders with general-purpose LLMs [22].

For instance, the BioReason [23] project successfully integrated a pretrained DNA model with an LLM by feeding sequence embeddings alongside textual questions into the model, achieving significant gains on tasks like pathway prediction and mutation analysis. This approach provides a valuable precedent for fusing biological sequence information with natural language reasoning.

### Virtual Screening Methodologies

In traditional small-molecule drug discovery, virtual screening involves several steps: scoring, ranking, docking, and hit selection [8], [24]. Scoring functions estimate binding affinity, ranking algorithms sort compounds, and docking predicts binding poses. Screening then selects high-scoring candidates from large libraries.

As a result, Ab-VS-Bench focuses on what can be inferred from sequence alone, without requiring structural models or docking. This makes our approach more similar to ligand-based virtual screening, where models infer binding potential from features rather than structures. Ab-VS-Bench bridges several research areas. It extends prior antibody modeling benchmarks by introducing a full virtual screening workflow; it enables instruction-following evaluation with LLMs; and it supports multimodal integration of protein representations into general-purpose language models. By doing so, it offers a new platform for testing interactive, language-driven antibody discovery at scale.

## III. Method

### A. Benchmark Tasks and Datasets

Ab-VS-Bench comprises three core tasks: Scoring, Ranking, and Screening, each supported by multiple datasets derived from experimental data on antibody or nanobody interactions. These datasets ensure biological relevance and have been processed into a unified format suitable for instructing large language models (LLMs). Table 1 summarizes the tasks, detailing their objectives, dataset examples, and evaluation metrics. Each task in Ab-VS-Bench includes datasets sourced from antibody experiments, divided into training, validation, and test sets where applicable. These splits follow standardized criteria from original sources or our stratified splitting methods. To prevent information leakage, test set sequences are excluded from any model variant’s training. For instance, in affinity maturation datasets adapted from AbBiBench [2], if a wild-type antibody or antigen was seen during pretraining, the specific mutant complexes in the test set were withheld from fine-tuning. We employ clustered splitting based on sequence similarity, as done in NbBench [7], to ensure test examples are novel, preventing models from merely memorizing training sequences. A key feature of Ab-VS-Bench is framing each task instance as a textual instruction-answer pair. We crafted prompts that clearly describe the problem to the model. Below are simplified examples of prompt formats for each task:

**TABLE I.**
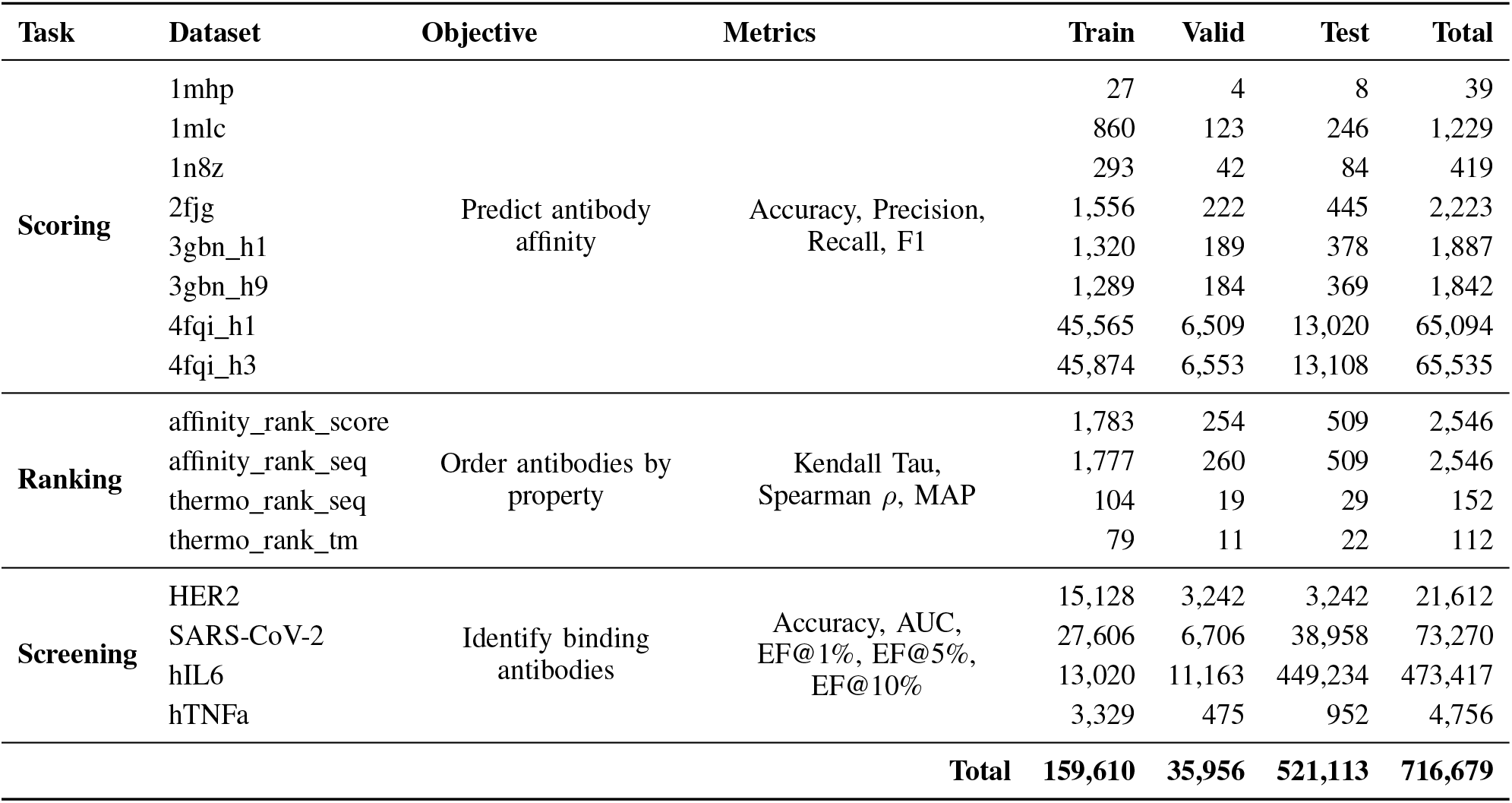
OVERVIEW OF AB-VS-BENCH: TASKS, OBJECTIVES, METRICS, DATASETS, AND DATA SPLIT STATISTICS.

| Task | Dataset | Objective | Metrics | Train | Valid | Test | Total |
| --- | --- | --- | --- | --- | --- | --- | --- |
| Scoring | 1mhp | Predict antibody affinity | Accuracy, Precision, Recall, F1 | 27 | 4 | 8 | 39 |
|  | 1mlc |  |  | 860 | 123 | 246 | 1,229 |
|  | 1n8z |  |  | 293 | 42 | 84 | 419 |
|  | 2fjg |  |  | 1,556 | 222 | 445 | 2,223 |
|  | 3gbn_h1 |  |  | 1,320 | 189 | 378 | 1,887 |
|  | 3gbn_h9 |  |  | 1,289 | 184 | 369 | 1,842 |
|  | 4fqi_h1 |  |  | 45,565 | 6,509 | 13,020 | 65,094 |
|  | 4fqi_h3 |  |  | 45,874 | 6,553 | 13,108 | 65,535 |
| Ranking | affinity_rank_score | Order antibodies by property | Kendall Tau, Spearman $\rho$, MAP | 1,783 | 254 | 509 | 2,546 |
|  | affinity_rank_seq |  |  | 1,777 | 260 | 509 | 2,546 |
|  | thermo_rank_seq |  |  | 104 | 19 | 29 | 152 |
|  | thermo_rank_tm |  |  | 79 | 11 | 22 | 112 |
| Screening | HER2 | Identify binding antibodies | Accuracy, AUC, EF@1%, EF@5%, EF@10% | 15,128 | 3,242 | 3,242 | 21,612 |
|  | SARS-CoV-2 |  |  | 27,606 | 6,706 | 38,958 | 73,270 |
|  | hIL6 |  |  | 13,020 | 11,163 | 449,234 | 473,417 |
|  | hTNFa |  |  | 3,329 | 475 | 952 | 4,756 |
| Total |  |  |  | 159,610 | 35,956 | 521,113 | 716,679 |

- **Scoring Prompt Example:** “Antigen: spike protein (RBD fragment); Candidate antibody sequence: EVQLVESGGGLVQPGGSLRLSCA… ; Question: Does this antibody bind the antigen strongly? (Answer ‘Yes’ or ‘No’)” – The model should respond with “Yes” or “No”. In regression cases, we might ask: “… Predict the binding affinity*K*_*d*_ of this antibody–antigen pair.” The model would output a numeric estimate or a category like “High affinity”.
- **Ranking Prompt Example:** “You are given 5 antibody variants targeting lysozyme, labeled A, B, C, D, E (sequences provided). Rank these antibodies from highest to lowest binding affinity to lysozyme.” – The model should output an order, e.g., C *>* A *>* B *>* E *>* D, possibly with justification. For thermostability: “Rank the following nanobodies by their melting temperature (higher Tm means more stable): 1, 2, 3, 4.”
- **Screening Prompt Example:** “Given the antigen = EGFR peptide and a library of 100 antibody clones (sequences 1…100), identify which clones are potential high-affinity binders.” – Due to prompt length constraints, we might split the library and query the model iteratively or ask for top-N candidates. The model might respond with a list of candidate IDs (e.g., “Clones 17, 42, 88 are likely binders”). Alternatively, we pose multiple-choice questions, e.g., “Out of antibodies A, B, C, which is most likely to bind the target?” and iterate.

In constructing these instructions, we transformed raw data (numbers, labels) into language. For instance, a binding affinity value is not presented as a number; instead, the prompt might say “antibody X binds with an IC50 of 50 nM (high affinity)” in a few-shot demonstration. We experimented with different phrasing templates and found that models often prefer explicit questions (e.g., “Which antibody is the best binder?”) and simple answers, versus open-ended generation. We standardized the expected answer format for each task (classification tasks expect a classification label; ranking tasks expect a sorted list, etc.), facilitating straightforward automatic evaluation by comparing the model’s output to the reference answer. We avoided direct numeric regression prompts, as initial trials showed LLMs struggled with precise numbers, often outputting generic sentences or uncalibrated estimates. Instead, we frequently reduced regression to discrete ordinal categories or relative comparisons, aligning with LLMs’ strengths in comparative reasoning and classification. Where numeric prediction is unavoidable, we accept a range of correct outputs.

Each task is converted into natural language instructions. For example, scoring may ask: “Given this antibody and antigen sequence, does it bind strongly?”; ranking may request: “Rank these 5 antibodies by binding affinity.”; and screening may prompt: “Which of these candidates are likely highaffinity binders?”

### B. Models

We experiment with multiple modeling approaches to evaluate the effect of domain-specific sequence knowledge and natural language understanding on antibody-related tasks. All examples are formatted using an *instruction tuning* format, allowing large language models (LLMs) to process task prompts and generate responses in a supervised manner. Where possible, regression outputs are discretized into categorical tiers to stabilize training.

#### a) Pretrained LLM (Zero-shot)

As a baseline, we evaluate general-purpose LLMs such as Mistral-7B and Qwen2.5-7B in a zero-shot setting, without any task-specific tuning. These models are prompted using plain-language instructions and evaluated directly. Although not trained for antibody-related tasks, they provide insight into whether general reasoning and language understanding can transfer to biological prediction tasks.

#### b) Instruction-Finetuned LLM

We fine-tune LLMs on task-specific instruction data. Each training example includes a natural language prompt and the expected output. This setup allows the model to learn mappings between textual descriptions and desired answers (e.g., scores, labels). Fine-tuning is conducted using a low learning rate with validation-based early stopping to prevent overfitting. This setup improves performance significantly over zero-shot LLMs and helps models better associate antibody properties with expected outcomes.

#### c) LLM with Antibody Embeddings (Multimodal, Cross-Attention Fusion [25])

Our primary model integrates antibody sequence information into the LLM using a cross-attention-based fusion strategy. As illustrated in Figure 1, we encode antibody sequences using a pretrained protein language model (e.g., ESM2 [15]), which remains frozen during training. The resulting embeddings are aligned with the LLM hidden states through cross-attention layers inserted into the LLM architecture.

**Fig. 1.**
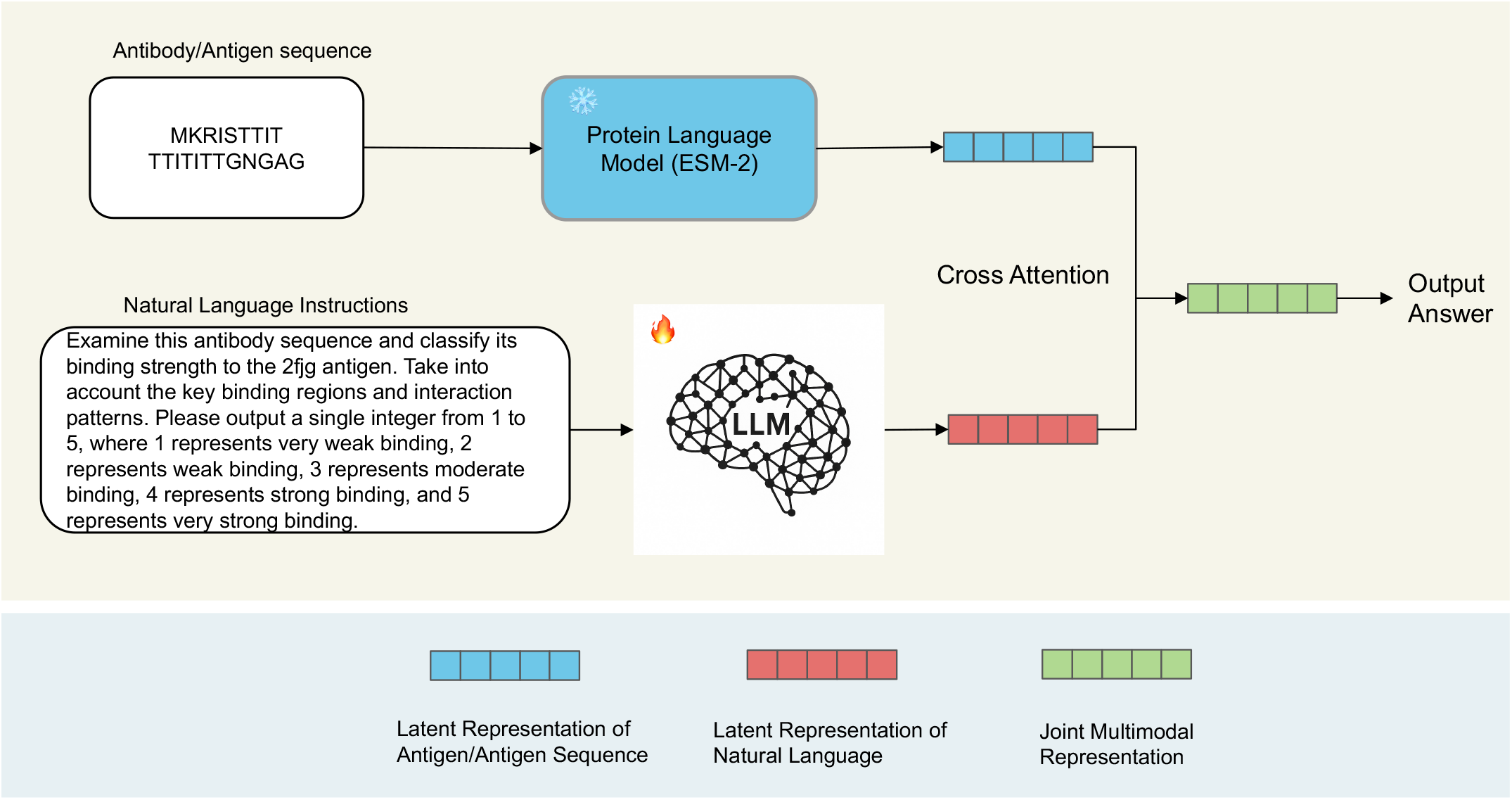
Overview of the proposed multimodal framework for antibody screening. The protein (antibody/antigen) sequence is first processed by a protein language model (ESM-2) to obtain a latent representation, while natural language instructions are encoded using a large language model (LLM). The two modalities are fused through a cross-attention module to form a joint multimodal representation, which is then used to generate the final prediction. The bottom legend illustrates the latent representations of each modality and their fusion.

During training, each input consists of a natural language instruction and the corresponding antibody sequence. The antibody is embedded independently and provided as an auxiliary input to the LLM. The LLM attends to both the tokenized prompt and the embedded antibody vector, allowing it to jointly reason over text and protein features. The antibody embeddings are injected at every decoder layer via a dedicated cross-attention module, enabling flexible information flow without modifying the LLM vocabulary.

To support stable training and maintain semantic alignment, we optionally enable Rotary Positional Encoding (RoPE) [26] on the embedded features. The overall model is fine-tuned end-to-end using the Hugging Face `trl` framework, with LoRA [27] applied to reduce memory cost and enable efficient adaptation.

#### d) Training Setup

All models are trained for 10 epochs using 4-bit quantization combined with LoRA adapters applied to key transformer modules. We adopt a batch size of 4 with gradient accumulation to achieve an effective batch size of 16, and we use a learning rate of 2 *×* 10^*−*4^. Early stopping is applied based on validation loss to prevent overfitting. For multimodal variants, cross-attention layers are employed to integrate antibody embeddings, with positional encodings added and the antibody encoder backbone kept frozen. This configuration allows the models to remain computationally efficient and interpretable, while effectively leveraging domain-specific sequence representations.

### C. Evaluation Metrics

Each task in Ab-VS-Bench is evaluated using metrics appropriate to its objective:

- **Scoring Tasks** aim to predict antibody binding affinity as a classification problem. For each dataset, we report standard classification metrics: Accuracy, Precision, Recall, and F1 Score. These metrics reflect the correctness and balance of predictions in distinguishing binding from non-binding antibodies.
- **Ranking Tasks** require models to order antibodies according to properties such as binding strength or thermostability. We use Kendall Tau [28], Spearman’s ***ρ***, and Mean Average Precision (MAP) to assess the quality of the predicted rankings. These metrics capture both the relative ordering and concentration of relevant items.
- **Screening Tasks** focus on early retrieval of true binders from a large candidate pool. We evaluate performance using Accuracy, Area Under the ROC Curve (AUC), and Enrichment Factors (EF) [29] at 1%, 5%, and 10%. EF metrics are particularly useful for assessing model utility in high-throughput screening scenarios, where only the top-ranked candidates matter.

All metrics are computed on the test sets, using the validation sets for model selection when available. The choice of metrics ensures a consistent and interpretable evaluation across tasks of varying difficulty and format.

## IV. Results

We evaluate our models across three categories of tasks: screening, scoring, and ranking. The comparison includes baseline models (direct and fine-tuned variants) and our proposed multimodal Mistral-Nemo model.

### A. Screening Tasks

Table II and Fig 2 together paint a clear picture of the screening benchmark. Across all four targets—HER2, SARS– CoV–2, hIL6, and hTNFa—the five evaluated LLM variants achieve nearly identical numbers: AUC stalls at 0.50 and enrichment factors hover around 1.0, while accuracy ranges from 0.48 to 0.96. The apparent paradox of “high” accuracy but chance-level AUC occurs because the datasets are profoundly imbalanced; Fig. 2 shows that positive binders constitute only a tiny fraction of each split, especially for hIL6 and hTNFa. Faced with such skew, every model collapses to predicting the majority class (non-binder) for all inputs. This strategy inflates accuracy—most examples really are negatives—yet fails to retrieve true binders, leaving AUC and EF no better than random. Consequently, the current task configuration masks performance differences among zero-shot, fine-tuned, and multimodal systems and provides little insight into a model’s genuine hit-discovery capability.

**TABLE II.** PERFORMANCE ON SCREENING TASKS. EACH METRIC’S BEST VALUE IS HIGHLIGHTED IN BOLD.

| Model | HER2 |  |  |  |  | SARS-CoV-2 |  |  |  |  | hIL6 |  |  |  |  | hTNFa |  |  |  |  |
| --- | --- | --- | --- | --- | --- | --- | --- | --- | --- | --- | --- | --- | --- | --- | --- | --- | --- | --- | --- | --- |
|  | Acc | AUC | EF(1%) | EF(5%) | EF(10%) | Acc | AUC | EF(1%) | EF(5%) | EF(10%) | Acc | AUC | EF(1%) | EF(5%) | EF(10%) | Acc | AUC | EF(1%) | EF(5%) | EF(10%) |
| Mistral-Nemo (Zero-shot) | 0.478 | 0.500 | 0.958 | 1.005 | 1.029 | 0.716 | 0.500 | 3.412 | 1.474 | 1.113 | 0.960 | 0.500 | 0.034 | 0.486 | 1.283 | 0.910 | 0.500 | 0.000 | 0.707 | 0.932 |
| Qwen2.5-7B (Zero-shot) | 0.478 | 0.500 | 0.958 | 1.005 | 1.029 | 0.716 | 0.500 | 3.412 | 1.474 | 1.113 | 0.960 | 0.500 | 0.034 | 0.486 | 1.283 | 0.910 | 0.500 | 0.000 | 0.707 | 0.932 |
| Mistral-Nemo (Fine-tuned) | 0.478 | 0.500 | 0.958 | 1.005 | 1.029 | 0.716 | 0.500 | 3.412 | 1.474 | 1.113 | 0.960 | 0.500 | 0.034 | 0.486 | 1.283 | 0.910 | 0.500 | 0.000 | 0.707 | 0.932 |
| Qwen2.5-7B (Fine-tuned) | 0.478 | 0.500 | 0.958 | 1.005 | 1.029 | 0.716 | 0.500 | 3.412 | 1.474 | 1.113 | 0.960 | 0.500 | 0.034 | 0.486 | 1.283 | 0.910 | 0.500 | 0.000 | 0.707 | 0.932 |
| Mistral-Nemo (Multimodal) | 0.478 | 0.500 | 0.958 | 1.005 | 1.029 | 0.716 | 0.500 | 3.412 | 1.474 | 1.113 | 0.960 | 0.500 | 0.034 | 0.486 | 1.283 | 0.910 | 0.500 | 0.000 | 0.707 | 0.932 |

**Fig. 2.**
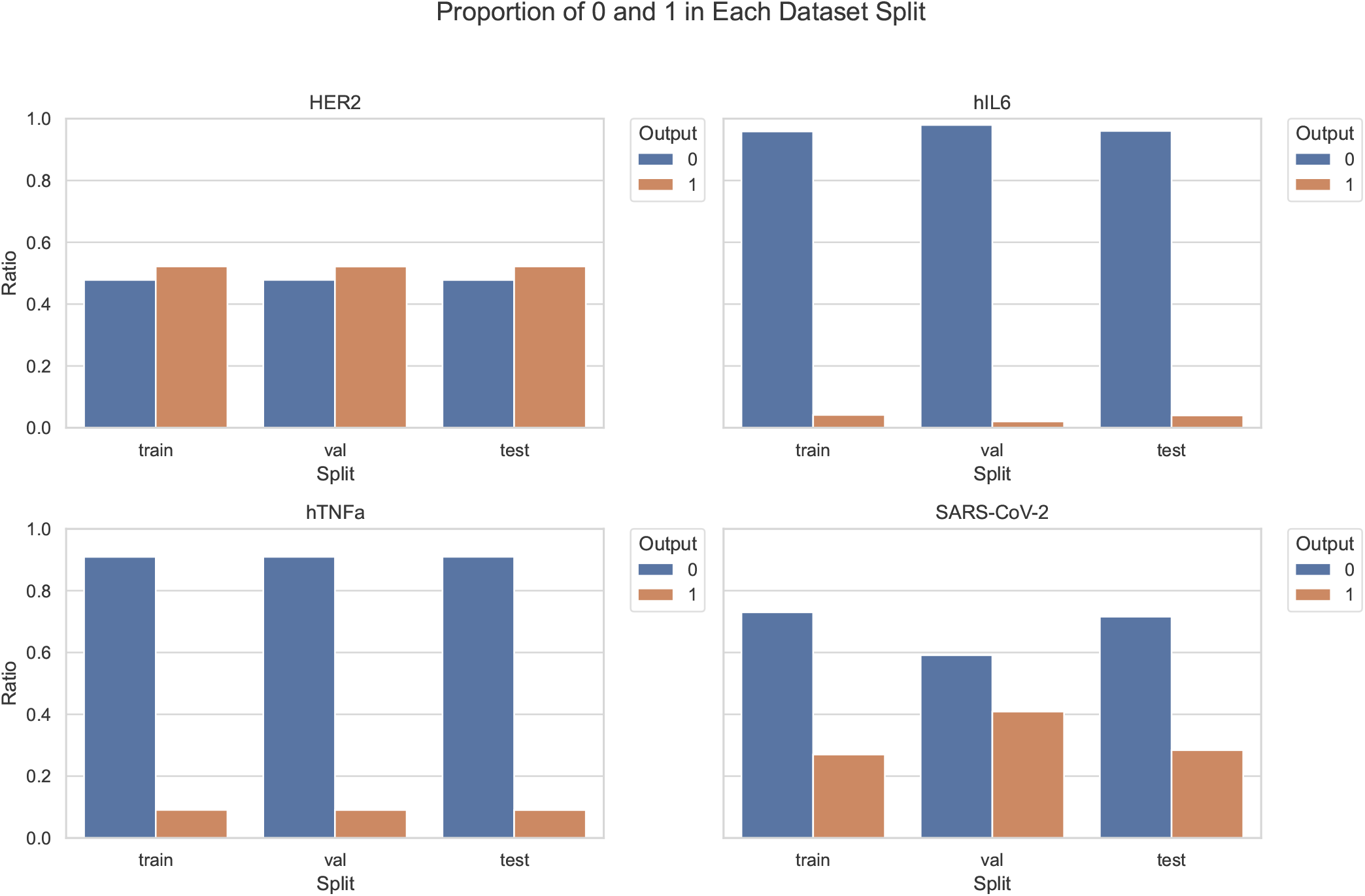
Proportion of positive (1) and negative (0) labels in the training, validation, and test splits for each dataset (HER2, hIL6, hTNFa, and SARS-CoV-2). The figure shows the class distribution across splits, illustrating varying degrees of class imbalance in different datasets.

To obtain a more discriminative benchmark, future work should incorporate training signals that emphasise rare positives, for example through balanced curricula, positive-class mining, or synthetic oversampling. Prompt or loss-function designs that penalise missed binders could further help LLMs overcome majority-class bias, and integrating structure-or similarity-based priors might steer generation toward plausible candidates before classification. Finally, evaluation protocols themselves need revision: metrics conditioned on a fixed number of generated molecules or balanced bootstrapped test sets would offer a more faithful measure of antibody discovery potential in low-prevalence settings.

### B. Scoring Tasks

Table III presents the performance of models on scoring tasks across eight antibody–antigen targets. Evaluation metrics include Accuracy, Precision, Recall, and F1 Score. Notably, fine-tuning on domain-specific scoring tasks leads to substantial performance gains. Both Mistral-Nemo (Fine-tuned) and Qwen2.5-7B (Fine-tuned) consistently outperform their zero-shot or direct counterparts, confirming the importance of task adaptation for LLMs in protein-related classification.

**TABLE III.**
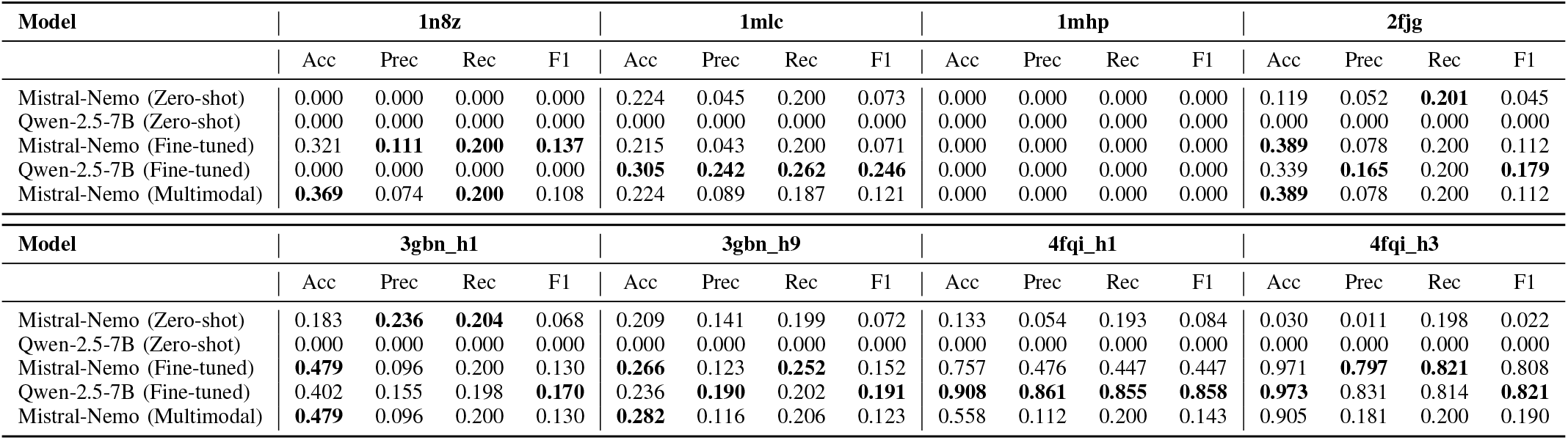
PERFORMANCE ON SCORING TASKS. EACH TASK REPORTS ACCURACY, PRECISION, RECALL, AND F1 SCORE. THE BEST VALUE FOR EACH METRIC IS IN **BOLD**.

The multimodal Mistral-Nemo model, which integrates antibody embeddings into the language model input, maintains competitive performance—particularly on the large-scale datasets 4fqi h1 and 4fqi h3. These datasets have tens of thousands of examples, enabling the model to benefit from sequence-level representation learning. However, despite its architectural sophistication, Mistral-Nemo does not consistently outperform the simpler fine-tuned LLMs. In several datasets, the fine-tuned Qwen2.5-7B even achieves higher F1 scores, suggesting that the integration of protein embeddings alone may not be sufficient to guarantee better downstream task performance.

This observation points to a critical insight: multimodality in itself is not a silver bullet. The effectiveness of integrating sequence embeddings depends heavily on alignment between modalities, the quality of the embeddings, and the way they are fused into the LLM. Future work should investigate improved fusion strategies or explore joint training of sequence encoders and LLMs to enable more cross-modality reasoning. Additionally, task-specific prompting or intermediate instruction tuning may offer lightweight alternatives to full fine-tuning or complex multimodal pipelines.

### C. Ranking Tasks

Table IV summarizes model performance on ranking tasks, using Kendall Tau, Spearman Rho, and MAP. Fine-tuned models outperform direct models by a large margin. Among all, the multimodal Mistral-Nemo model achieves the best ranking performance, especially on the Affinity Rank Score task (Kendall = 0.994, Spearman = 0.996, MAP = 0.997), indicating its effectiveness in capturing fine-grained differences in antibody ranking.

**TABLE IV.** PERFORMANCE ON RANKING TASKS. EACH TASK REPORTS KENDALL TAU, SPEARMAN RHO, AND MAP METRICS.

| Model | Affinity Rank Score |  |  | Affinity Rank Seq |  |  | Thermo Rank Seq |  |  | Thermo Rank Tm |  |  |
| --- | --- | --- | --- | --- | --- | --- | --- | --- | --- | --- | --- | --- |
|  | Kendall | Spearman | MAP | Kendall | Spearman | MAP | Kendall | Spearman | MAP | Kendall | Spearman | MAP |
| Mistral-Nemo (Zero-shot) | -0.005 | 0.015 | 0.232 | 0.096 | 0.094 | 0.372 | -0.050 | -0.059 | 0.403 | 0.147 | 0.197 | 0.374 |
| Qwen2.5-7B (Zero-shot) | -0.014 | -0.002 | 0.171 | 0.102 | 0.107 | 0.416 | -0.028 | -0.045 | 0.335 | 0.186 | 0.229 | 0.356 |
| Mistral-Nemo (Fine-tuned) | 0.962 | 0.967 | 0.986 | 0.928 | 0.935 | 0.977 | 0.531 | 0.552 | 0.742 | **0.545** | **0.605** | **0.826** |
| Qwen2.5-7B (Fine-tuned) | 0.516 | 0.548 | 0.852 | 0.555 | 0.589 | 0.848 | 0.322 | 0.353 | 0.746 | 0.416 | 0.498 | 0.648 |
| **Mistral-Nemo (Multimodal)** | **0.994** | **0.996** | **0.997** | **0.942** | **0.949** | **0.978** | **0.559** | **0.614** | **0.806** | 0.473 | 0.491 | 0.790 |

### D. Cross-Target Generalization

To assess generalizability, we conduct transfer experiments by training models on the large-scale datasets 4fqi h1 and 4fqi h3, then evaluating on six unseen antibody–antigen targets. As shown in Table V, models trained on 4fqi h3 consistently outperform those trained on 4fqi h1 across most targets. This suggests that 4fqi h3 provides broader coverage or more informative patterns, making it a more effective source for cross-target transfer.

**TABLE V.**
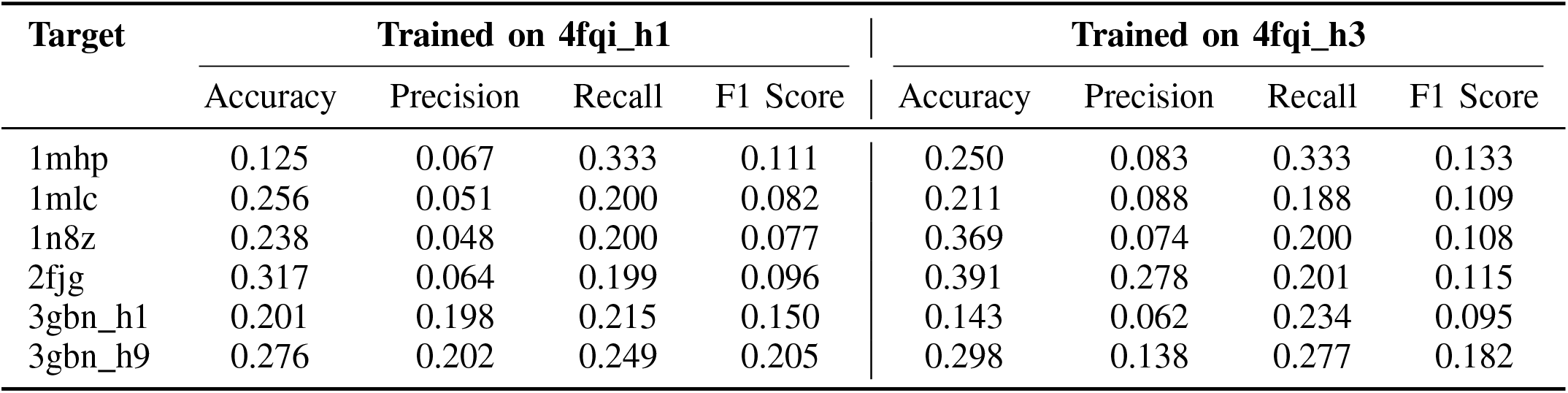
PERFORMANCE COMPARISON ON DIFFERENT TARGETS, TRAINED ON 4FQI H1 AND 4FQI H3.

These results highlight a potential synergy effect in fine-tuned LLMs—once exposed to a sufficiently large and diverse training set, the model appears to internalize generalizable features that transfer well to new antibody–antigen pairs. This insight is especially valuable for real-world antibody discovery, where training data is often limited for emerging or rare targets. In such low-resource scenarios, it may be beneficial to fine-tune LLMs on large datasets from related targets and apply them zero-shot or few-shot to new ones. Future work could explore target similarity metrics or pretraining curriculum strategies to maximize this transfer potential.

## V. Conclusion

We introduce Ab-VS-Bench, a novel benchmark for evaluating large language models on virtual antibody screening tasks through natural language instructions. Covering scoring, ranking, and screening tasks, our benchmark enables the first systematic assessment of direct, fine-tuned, and multimodal LLMs in this domain. Experimental results show that fine-tuning greatly enhances performance, especially under data scarcity, while the multimodal Mistral-Nemo model achieves strong generalization. However, challenges such as class imbalance and modest gains from multimodal integration highlight room for improvement. Ab-VS-Bench provides a foundation for developing more robust, instruction-following models for antibody discovery.

## Footnotes

1 The dataset is publicly available at: https://huggingface.co/datasets/ZYMScott/Ab-VS

## Notes

### Competing Interest Statement

The authors have declared no competing interest.

